# Cellular feedback to organotelluranes displays modulation of antioxidant proteins gene expression

**DOI:** 10.1101/2021.01.05.425411

**Authors:** Felipe S. Pessoto, César H. Yokomizo, Rodrigo L. O. R. Cunha, Iseli L. Nantes-Cardoso

## Abstract

Organotelluranes RT3 and RT4 are thiol reagents that induce mitochondrial transition pore (MTP) opening in a sensitive and insensitive manner to cyclosporin A. Although RT3 and RT4 promote glutathione depletion, paradoxically, they are also an efficient antioxidant for membrane lipids. These compounds’ antagonistic effects elicited the challenging question of how the gene expression of antioxidant enzymes would respond to treatment with these compounds. The influence of RT3 and RT4 on antioxidant enzyme expression was investigated in cultured aortic smooth muscle cells (ASMC). RT3 and RT4 promoted disruption of ionic calcium homeostasis, mitochondrial transmembrane potential (ΔΨ), and cell death in a dose-dependent manner. The cell death mechanisms responded qualitatively to the increase of the organotellurane concentration and changed from apoptosis to necrosis. RT3 and RT4 increased the expression of thioredoxin significantly. RT3 also increased the expression of glutaredoxin and glutathione peroxidase, slightly the catalase expression without significant effects on SOD expression. The results are consistent with GSH and protein thiol depletion and discussed based on the cell toxicity mechanism exhibited by these compounds.

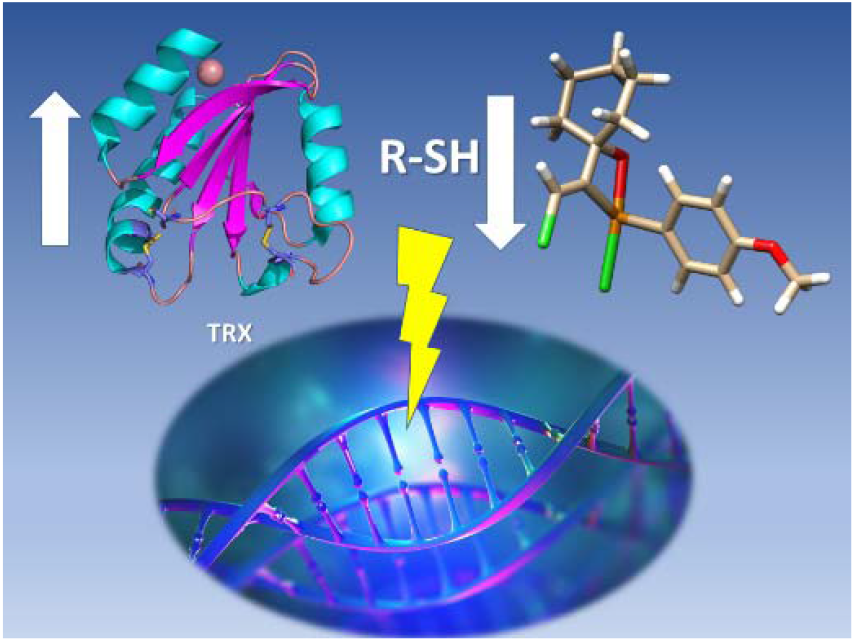

## Introduction

Tellurium forms inorganic and organic derivatives, both with a multiplicity of biological effects that are attractive due to the potential therapeutic applications and technological applications.(Comasseto et al. 2007; Cunha et al. 2009; Hagar et al. 2020; Machado and Pereira da Silva 2020; Medina-Cruz et al. 2020; Pietrasiak and Togni 2017; Srivastava et al. 2018; Teixeira et al. 2018; Moroder and Musiol, 2019; Sato et al. 2019; Wang et al. 2020) The organic derivatives of tellurium, are compounds with at least, one tellurium-carbon ligation in their structures.(Cunha et al. 2009; Nogueira et al. 2004; Pietrasiak and Togni 2017; Teixeira et al. 2018) Organotellurium comprise the groups of the divalent derivatives tellurols, diorganoditellurides, and diorganotellurides, and the hypervalent derivatives that are the organotellurium trihalides, diorganotellurium dihalides, organotellurium oxides, organotellurates, and organopertelluranes.(Cunha et al. 2009) Most of the biological effects of tellurium compounds are related to the reactivity with thiol groups.(Cunha et al. 2009; Kheirabadi and Izadyar 2020; Nantes et al. 2011; Pessoto et al. 2007; Yokomizo et al. 2015) The inorganic tellurium compounds AS-101 and SAS form stable Te-Cys_4_ complexes by replacing of labile ligands by four equivalents of cysteine.(Cunha et al. 2009; Silberman et al. 2016) Silberman et al demonstrated that distinct of inorganic tellurium compounds, the diaromatic organotelluranes of the type Te^IV^Ar_2_X_n_ (X = Cl, O and CH_3_COO) do not form stable complexes with cysteine.(Silberman et al. 2016; Teixeira et al. 2018) The reaction mechanism of diaromatic organotelluranes forms complexes with two cysteines that are intermediates of the final products, the Te^II^Ar_2_ and cystine.(Silberman et al. 2016) The effects of trichlorotelluro-dypnones on the bioenergetics of cells and their relationship with thiol reactivity were also investigated.(Yokomizo et al. 2015) For these compounds, it was proposed a reaction mechanism like to SN2-type and SN1-type that are associative and dissociative mechanism.(Yokomizo et al. 2015) All the postulated mechanisms implicate in the depletion of thiol groups with repercussions on the cell redox balance.(Allen and Mieyal 2012; Benhar 2020; Jones 2006; McBean et al. 2015; Paul et al. 2018) GSH is considered the most abundant redox buffer in cells and acts in conjunction with proteins, particularly the TRX system (TRX, TRX reductase, and NADPH) in maintaining the intracellular redox homeostasis.(Arnér and Holmgren 2000; Berndt et al. 2006; Bjørklund et al. 2020; Jones 2006; McBean 2017; Panday et al. 2020) The cells have redox-sensitive transcription factors that regulate gene expression in response to changes in the cellular redox status. The pathway of redox-sensitive gene expression involves agents acting as sensors, transducers and effectors.(Gong et al. 2020; Hirota et al. 2000; Spector et al. 1988; Messina et al. 2019) The redox state of cysteine residues is a key signal for a cellular gene expression in response to loss of redox homeostasis. In this regard, distinct oxidative states of cysteine residues elicit specific responses in cells. The reactivity of thiol groups depends on their protonation state. The thiolate group (−S^−^) is more reactive with its conjugate acid form (−SH).(Castranova et al. 2016; Wall et al. 2012) Also, the oxidation of protein bound thiol groups depends on the accessibility to the oxidative agents and different derivatives can be formed, i.e. sulfenic (−SOH), sulfinic (SO_2_H) and sulfonic (SO_3_H) acids.(Icimoto et al. 2017; Jeong et al. 2011) The sulfenic and sulfinic groups are reversible oxidized states of thiol group while sulfonic acids represent irreversible modification of proteins. The thiol group of proteins are also targeted by the nitrosative species nitric oxide (NO^•^) and peroxynitrite (ONOO^−^) that is formed by the reaction of NO^•^ with superoxide ion (O_2_^−•^). The toxicity of nitrosative species results of covalent binding of NO^•^ to the thiol group of proteins leading to the formation of S-nitrosothiols (SNOs)(Sun et al. 2006) and the occurrence of protein nitration by the appending of nitro group (−NO_2_) to a tyrosine residue.(Paes de Barros et al. 2020) A diversity of enzymes and transcription factors responds to the oxidative stress such as peroxiredoxins,(Jeong et al. 2006; Jiang et al. 2020; Kang et al. 2004; P. et al. 2017; Rhee et al. 2005) the transcription factors NF-κB and AP-1,(Attafi et al. 2020; Gius et al. 1999; KLATT et al. 1999; Nishi et al. 2002) the negative regulator of Nrf2, protein 1 (Keap1),(Priya Dharshini et al. 2020; Wakabayashi et al. 2004) thioredoxin(Holmgren 1995; Rashidi et al. 2019; Shao et al. 2020; Sumbayev 2003; Zeng et al. 2020), tyrosine phosphatases (PTP),(Denu and Tanner 1998; Hsu et al. 2020) and the kinase MEKK1.(Cross et al. 2007; CROSS and TEMPLETON 2004; Matsushita et al. 2020) As an example, the transcription factor Nrf2 controls hundreds of genes involved in the combat of the oxidative stress. Nrf2 is repressed by association with the homodimeric form of the Keap1 protein. Human Keap1 has several of its 25 cysteine residues with low p*K*_a_ and consequently, they are highly reactive in the cell conditions. Therefore, Keap1 protein regulates Nrf2 in response to the cell redox state.(Baird and Yamamoto 2020; Janssen-Heininger et al. 2000; Krajka-Kuźniak et al. 2017; Limón-Pacheco and Gonsebatt 2009) The redox-sensitive gene expression is also regulated by S-glutathionylation, that is the reversible formation of disulfide bonds between GSH and cysteinyl residues.(Limón-Pacheco and Gonsebatt 2009; Mailloux et al. 2020; Matsui et al. 2020) The S-glutathionylation is a negative regulatory process for tyrosine hydroxylase,(Borges et al. 2002) PKC-α,(Ward et al. 2000) MEKK1 kinase,(Cross et al. 2007) PTP1B,(Lee et al. 1998; William C. Barrett et al. 1999) and inhibitor of kappa B kinase-β, which inhibits NF-κB(Checconi et al. 2019; Reynaert et al. 2006) that in turn is reversed by glutaredoxin (Grx).(Gallogly and Mieyal 2007; Wen et al. 2020) TRX is a low molecular weight disulfide oxido-reductase enzyme with 10-12 kDa. TRX has two reactive cysteine residues that forms a disulfide bond upon oxidation and that is recycled the action of TRX reductase and NADPH.(Collet and Messens 2010) TRX exerts its antioxidant activity by the enzymatic action on other enzymes and transcription factors as well as the scavenging of reactive oxygen species.(Berndt et al. 2006)(Collet and Messens 2010) TRX is overexpressed in conditions in which the redox-active cysteines are inactivated by heavy metals, and reactive species produced by ischemia–reperfusion and viral infection.(Branco et al. 2019; Kasuno et al. 2003; Zhou et al. 2020) In the present study, the toxicologic effects of RT-03 and RT-04 (Fig. 1) on aortic smooth muscle cells (ASMC) and the repercussions on the expression of thioredoxins (TRx), thioredoxin reductase, glutaredoxin, SOD, catalase, and glutathione peroxidase were investigated.

**Figure 1.**
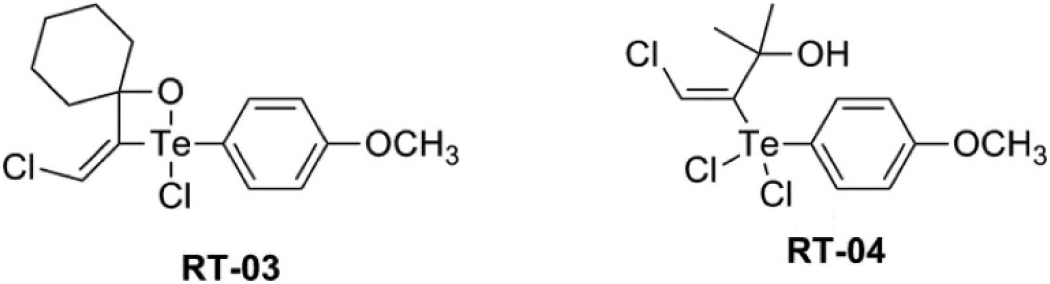
Molecular structures of RT-03 and RT-04.

## Materials and Methods

### Synthesis of organotelluranes

The organotelluranes used (RT-03 and RT-04) were synthesized by the electrophilic addition of p-methoxyphenyl tellurium trichloride to alkynes, called 1-ethynyl-1-cyclohexanol and 3-methyl-3-hydroxy-butyne, respectively.(Zeni et al. 1999)

### Animal handling and euthanasia

Rats of the Wistar lineage, maintained and euthanized using procedures approved by the internal ethics committee registered at COBEA, which supervises all norms and ethical principles in maintenance of animals, such as: space, ventilation, temperature, nutrition, hydration and proper handling. Only males of the Wistar lineage were used in this research, the euthanasia method used was the cervical dislocation for rats up to 200 grams, and cranial trauma for animals over 200 grams, these methods are described in Guide to the care and use of experimental animals, Canadian Council on Animal Care (CCAC).(Olfert, Cross and McWilliam, 2020) The chemical method of euthanasia cannot be used because in this type of study it is contraindicated due to the interaction of the lethal chemical agents with membranes, interfering in the transport of ions, among other effects.

### Isolation of hepatic mitochondria

Mitochondria were isolated from the liver of male Wistar rats of approximately 200 g using the differential centrifugation technique.(Schneider 1951) The liver was removed after the animal’s death and washed in a solution containing 250 mM sucrose, 10 mM HEPES-KOH buffer and 1 mM EGTA at pH 7.4 at 4 ° C, chopped into small fragments and homogenized in Potter-Elvehjen. The suspension was centrifuged at 580g for 5 minutes and the supernatant purchased was centrifuged at 10,300g for 10 minutes. The resulting supernatant was discarded along with the upper lipid phase with Pasteur pipette, and the resulting pellet resuspended in approximately 25 ml of medium containing 250 mM sucrose, 10 mM HEPES-KOH buffer and 0.3 mM EGTA at pH 7.4 at 4 °C and centrifuged at 3400g for 15 minutes. The final pellet or mitochondrial fraction was resuspended in a solution containing 250 mM sucrose, 10 mM HEPES-KOH buffer at pH 7.4 at 4 ° C, at a concentration of approximately 100 mg of protein per ml.

### Dosage of proteins

The protein concentration was quantified by the Biuret method, where an aliquot (10 mL) of the sample to be determined was mixed with 100 μL of a solution of 5% deoxycholic acid (m / v) and water q.s. 1.5 mL. To this mixture was added 1.5 mL of the biuret reagent, composed of 0.15% (m/v) cupric sulfate, 0.6% (w/v) sodium and potassium tartrate and 0.75 M NaOH. After 10 min, the absorbance was determined at 540 nm against a blank of reagents and a 1% BSA solution was used as standard.(Layne 1957)(Noble and Bailey 2009)

### ANT extraction of rat liver mitochondria

Isolation of ANT was done according to Rück et al, 1998.(Rück et al. 1998) All steps were done at 4°C. Isolated liver mitochondria (20 mg) in 1 ml of isolation medium were incubated for 5 min with an equal volume of extraction buffer consisting of 40 mM KH_2_PO_4_, 40 mM KCl, 2 mM EDTA and 6% Triton X-100, at pH 6.0. The suspension was centrifuged for 30 min at 24,000 g and the supernatant was loaded onto a column packed with 1g of dry hydroxyapatite (Bio-Rad). After elution with extraction buffer the fractions collected containing proteins were diluted in 20 mM MES buffer, 0.2 mM EDTA and 0.5% Triton X-100 at pH 6.0 in the ratio 1: 1. This sample was applied to a 1-ml cation exchange HiTrapSP column (Pharmacia), connected with a FPLC system and eluted by a NaCl gradient (0.1 M NaCl in column buffer). Fractions were analyzed by 12% SDS-PAGE and by immunodetection by anti-ANT, porin, CyPD, and Bax antibodies after transfer to nitrocellulose sheets.

### MALDI-TOF Spectrometry

The mass spectra were acquired in a MALDI-TOF Pro mass spectrometer device that can operate in linear or reflectron mode using harmonic reflecton which increases the resolution and narrows the focusing time. The samples were mixed with saturated solution (5 mg / ml) of 3,5-dimethoxy-4-hydroxycinnamic acid in 50% acetonitrile and 0.5% TFA, and 0.5 ml of the mixture was loaded onto the MALDI stainless steel films for analysis.(Kawai et al. 2005)

### Culture of rabbit aortic smooth muscle cells

Maintenance of rabbit aortic smooth muscle cell (ASMC) culture was performed every 3 days. The removal of the cells from the flasks, when confluent, was performed by the trypsinization process. After determining cell viability, 1 × 10^6^ cells were distributed in flasks of 75 cm^3^ and grown in DMEM medium (Sigma Chemical Co.) supplemented with 10% fetal bovine serum (BFS, Gibco, BRL-USA), and kept in a 5% of CO_2_ at 37°C. The trypsinization for cell handling was done by discarding of the culture medium followed by cell washing, at least 2 times with PBS solution. In sequence, a 2.5% trypsin solution containing 0.25% EDTA was added to the flask and held for 2 minutes at 37 °C to inactivate the reaction and then culture medium supplemented with BFS was added in a volume 2 times greater than that of trypsin used. After cells were detached from the flask, the solution was centrifuged at 450 g for 10 min at 20 °C and cell viability was determined.(Rodrigues et al. 2007)

### Cell viability assays

The yellow tetrazole MTT (3-(4,5-dimethylthiazolyl-2) −2, 5-diphenyltetrazol bromide) is metabolically reduced by living cells, in part by the action of dehydrogenase enzymes, to generate reduced equivalents such as NADH and NADPH. In the intracellular medium of viable cells, the reduction of the yellow tetrazole salt gives the purple formazan salt, which can be solubilized and quantified by spectroscopic measurements. The assay for EC50 determination was performed in 96-well plate, in each well, 2 × 10^5^ cells were added, at least 12 hours after cell distribution that is the minimum time for cells to adhere on the plate. After the incubation time had elapsed, 10 μl of the MTT solution (5mg/ml) (Invitrogen Co., USA) was added to each well and the plate incubated for an additional 4 hours. The whole procedure was performed in the dark due to photosensitivity of the MTT reagent. After the incubation time, 100 μl of a 10% SDS solution in 0.01 M HCL was added to each well, and after incubation for 9 to 12 hours the absorbances at 570 and 650 nm of each well was read using a reader of (Biotek ELX800). Blue trypan assay was performed on 25 × 13 mm petri dishes containing 5 × 10 5 cells. After incubation, 200 μl of the 0.4% tripan blue solution was added and nine fields / plate were analyzed, 50 cells / field counted, in a total of three plates per concentration. The percentage of dead cells was calculated in relation to the total cells present in the field. For the V-FITC/7-AAD assay, the cells were trypsinized and centrifuged 500 g for 5 min. The binding medium containing 10 mM of HEPES, 140 mM NaCl, 2.5 mM CaCl_2_, pH 7.4, and 1 μg/mL annexin V were added to the precipitate. Cells were incubated in light-protected propylene tubes and kept at room temperature for 15 min. Detection of the percentage of cells in apoptosis and necrosis was determined on a flow cytometer (Beckman Coulter, Lab Cell Quanta SC MPL model).

### Intracellular Ca^2+^ measurements

Rabbit aortic cells were loaded with 10 μM of Fura-2AM. Before starting the experiments, the cells were washed with Ca^2+^-free buffer to avoid interference of the external Ca^2+^. Ca^2+^ levels were measured on isolated intact cells using a high resolution inverted digital microscope that was coupled to a CCD camera controlled by a software.

### Intra-cell mitochondrial membrane potential

Rabbit aortic cells were loaded with 1 μM TMRM (tetramethyl rhodamine) for 30 min. Before starting the experiments, the cells were washed with fluorescence buffer. Fluorescence was monitored in isolated intact cells using a high-resolution digital microscope with an inverted microscope coupled to a CCD camera.

### Oxidation of total thiol groups

After induction of thiol group oxidation by the organotelluranes or 0.6 mM t-BOOH, the thiol content of cells was determined using 5,5’dithiobis (2-nitrobenzoic acid) (DTNB). The samples were centrifuged 2 min at 10,000 rpm and the pellet was treated twice with 200 μL of 6.5% trichloroacetic acid (TCA). After the addition of TCA, the samples were submitted to centrifugation at 10,000 g during 5 min to precipitate the protein content. The final pellet was resuspended in 1 mL of the cell medium in which 100 μM 5,5’-dithio-bis(2-nitrobenzoic acid) (DTNB), 0.5 mM EGTA, and 0.5 M Tris, pH 8.3 were present. The colorimetric determination of SH was measured by the absorbance intensity at 412 nm, using cysteine for calibration.(Castilho et al. 1996)

## Results and Discussion

### Effects of RT-03 and RT-04 on ASMC viability and bioenergetics

The ASMCs were incubated with organotelluranes RT-03 and RT-04 at the concentrations of 0, 10, 25, 50, 75, and 100 μM for one hour and the cell viability and death mechanisms were determined by flow cytometry using the annexin V-FITC/7-AAD kit. (Figure 2A and B).

**Figure 2.**
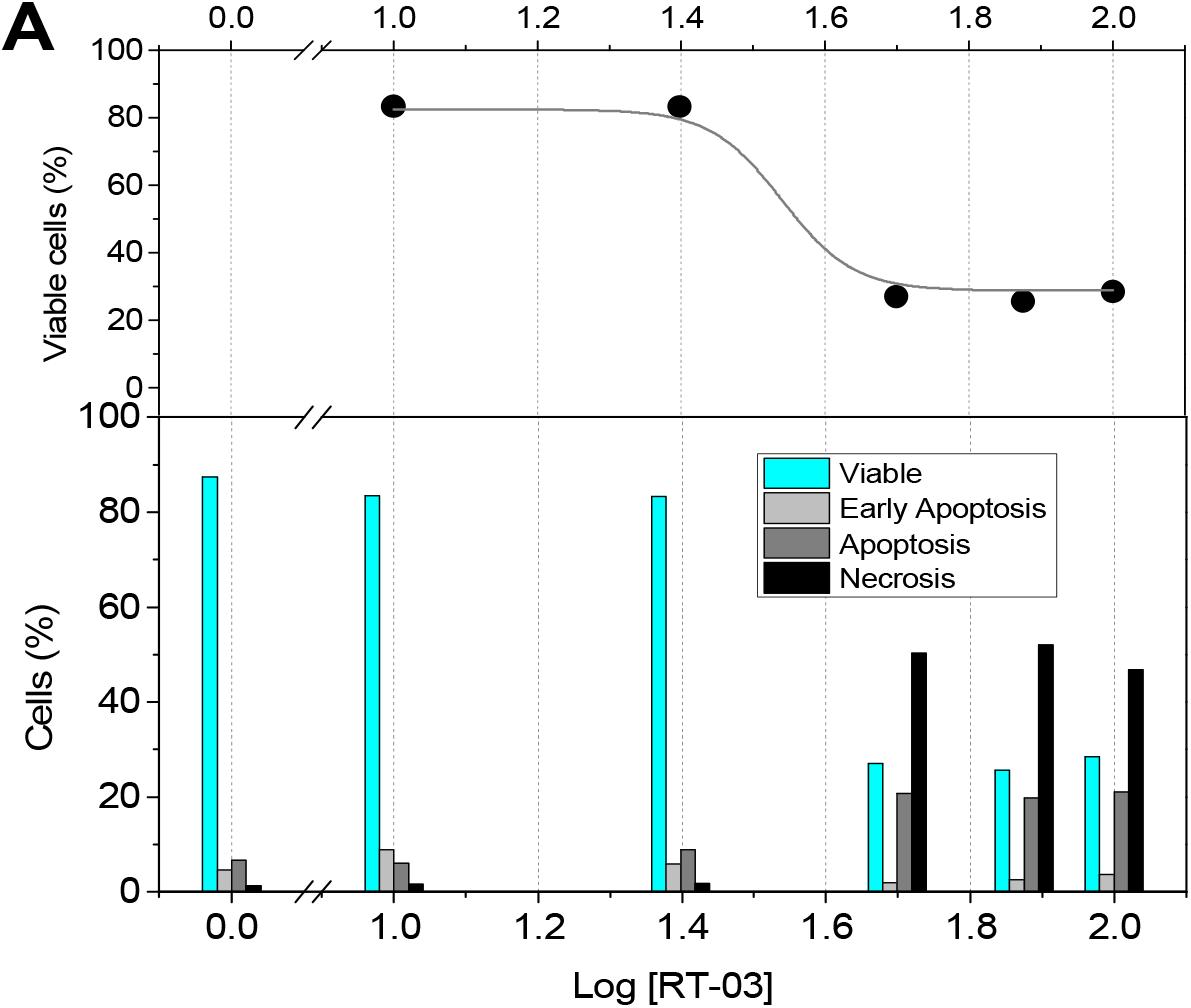
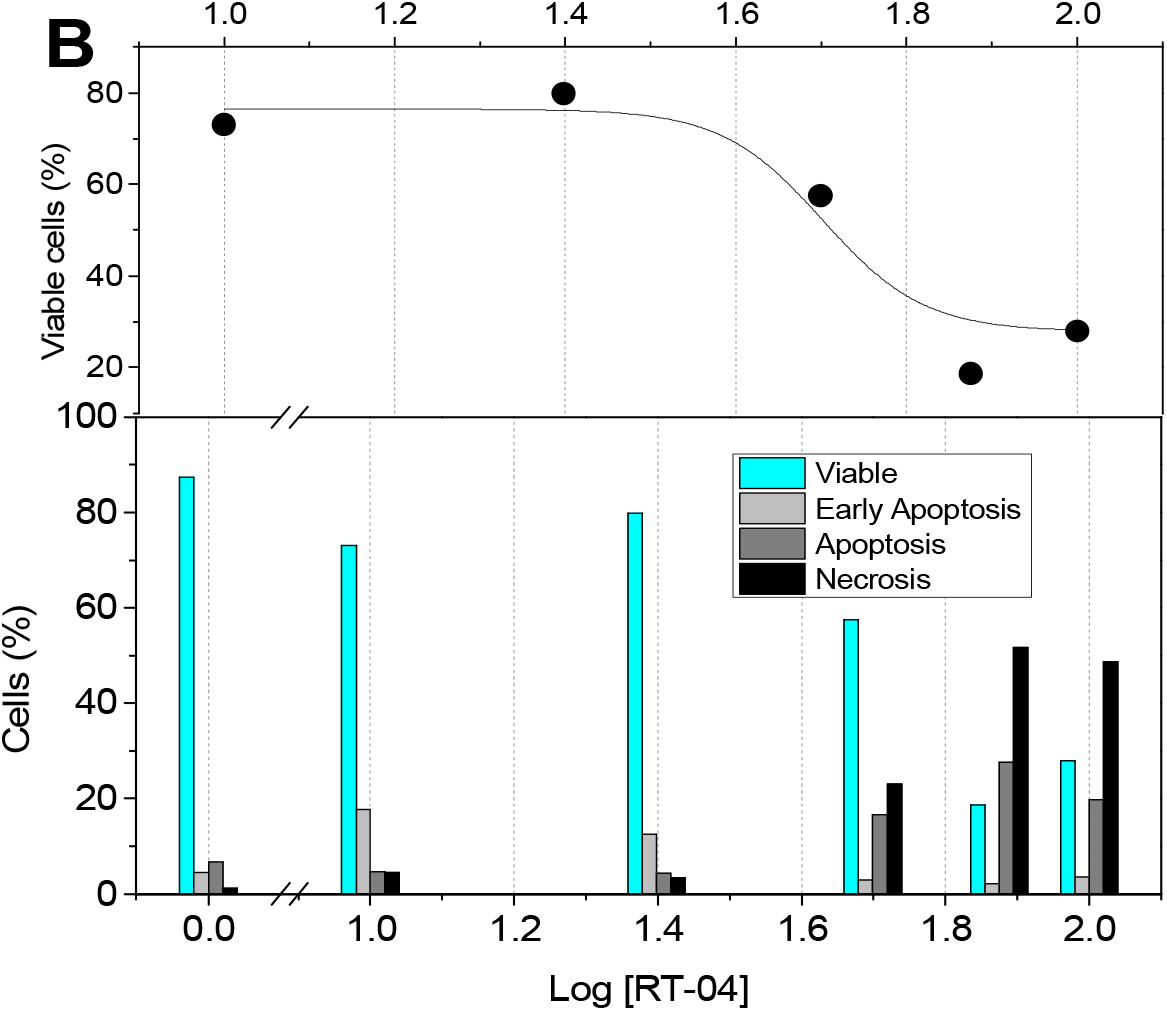
Flow cytometry of MLAC cells incubated with organotelluranes. MLAC cells (1×10^5^ cells) were incubated for 1h at 37 °C under 5% CO_2_ atmosphere, with RT-03 (panel A) and RT-04 (panel B). The insets show the curves for viable cells EC_50_ for each compound treatment. The percentages of apoptotic and necrotic cells were determined by flow cytometry using double labelling with annexin V-FITC and 7-AAD.

In viable cells, phosphatidylserine (PS) is restricted to the inner leaflet of the plasma membrane at expenses of ATP and it is not accessible for the specific staining with fluorescein-labeled annexin V. In the early step of apoptosis, the asymmetric PS distribution is randomized and makes the phospholipid accessible for labeling. During this phase, the membrane of dying cells is stained by fluorescein-labeled annexin V before the occurrence of DNA hydrolysis and lose of cellular morphology. Therefore, the annexin V staining indicates that cells are in the early apoptosis. The progression of apoptosis allows the staining of cells by the DNA-specific viability dye 7-AAD and characterizes late apoptotic cells.(Pessoto et al. 2015)(Schlegel and Williamson 2001)(Zachowski et al. 1989) Otherwise, the staining only with 7-AAD characterizes necrotic cells. Figures 2(a), and 2(b), lower panels, show the results of cell death mechanisms promoted by RT-03 and RT-04, respectively. The data of cell viability were used for the calculation of EC_50_ (Figures 2, upper panels). Figure 2A and B, upper panels, show the percentage of viable cells as a function of the logarithm of organotellurane concentration. In this condition, the EC_50_ for cell viability was calculated by fitting the data by equation 1.

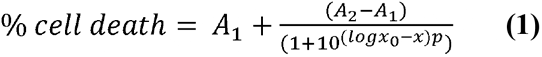

Where, *A*_1_ and, *A*_2_ are respectively the upper and lower asymptotic of the curve, *p* (Hill slope) the tangent slope to the curve and EC_50_ calculation from the curve turning point (*logx*_0_). The EC_50_ determined for RT-03 e RT-04, were, respectively = 34±4 and 50±5 μM. Cell viability was also determined using Trypan blue assay (not shown). The same EC_50_ for RT-03 (34±3 μM) was obtained using Trypan blue. The EC50 of RT-04 determined by Trypan blue assay was 64±3 μM and is close of the value obtained from flux cytometry data (50±5 μM). These results show the correlation between the events measured by both techniques. Therefore, the uptake of telluranes by the cells produces a toxic effect that triggers the mechanisms of programmed cell death whose events that signal its occurrence were measured by Annexin V / 7-AAD. At the same time these events lead to permeabilization of the plasma membrane which is measured by the uptake of Tripan blue. It is important to note that the significantly lower EC_50_ of RT-03 in relation to RT-04 (32-47%, considering Annexin V / 7-AAD and trypan blue, respectively) is consistent with the results previously obtained in isolated rat liver mitochondria (IRLM).(Yokomizo et al. 2015) In IRLM, RT-03 was also more efficient than RT-04 to promote damages in the organelle. The treatment of ASMCs with RT-03 at concentrations of 10-25 μM increased the percentage of apoptosis and necrosis by 31% and 44% respectively over the control. The use of 10-25 μM of RT-04 increased apoptosis up to 50% and necrosis up to 164%. Although the percentage of apoptosis had increased significantly after the treatment with 25 μM of the organotelluranes, around 75% of ASMCs remained viable. Bellow EC50 apoptosis responded predominantly for the cell death and above the EC_50_, necrosis was the predominant cell death mechanism promoted by the organotelluranes. The cell viability EC_50_ values of RT-03 and RT-04 were close to the EC_50_ of SH depletion that were, respectively, 60.2 ± 3 and 68.7 ± 3 μM (Figure 3). Figure 3 shows also that RT-03 and RT-04 did not deplete GSH of ASMCs. These results suggest that cell death promoted by the organotelluranes are related to the depletion of protein thiol groups but not to the oxidative stress. In IRLM, RT-03 and RT-04 promoted depletion of both protein thiol and GSH. The concentration of mitochondrial and cytosolic GSH are both in the range of 10-15 mM.(Marí et al. 2009) However, the enzymatic machinery to synthesize GSH is present only in cytosol as well as the pentose-phosphate pathway that is an important source of NADPH (48). The organotelluranes, RT-03 and RT-03 react as Lewis acids leading to protein SH depletion according to two possible mechanisms that were previously postulated: bis alkylation and redox mechanism.(Pessoto et al. 2007)

**Figure 3.**
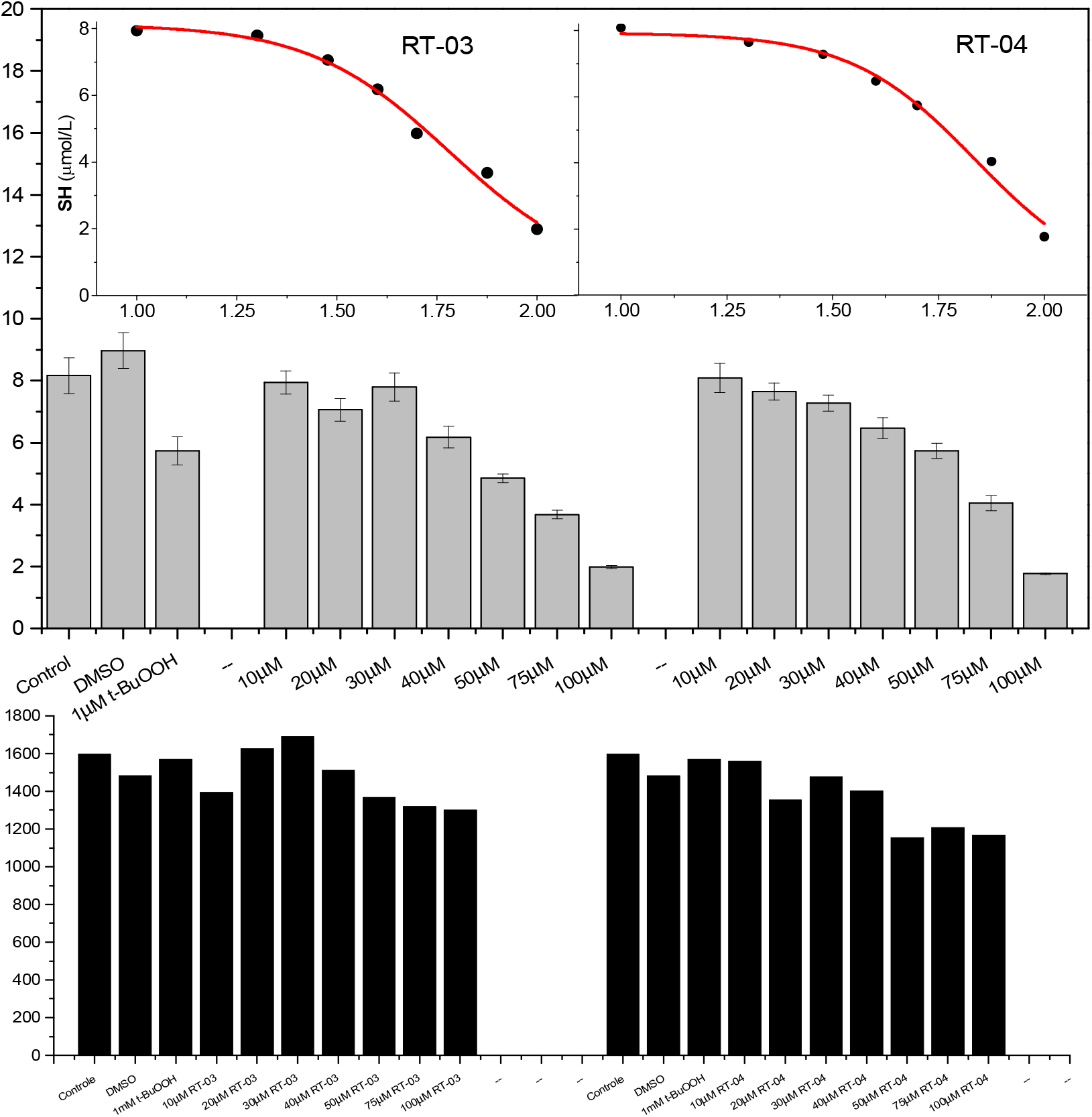
Effect of RT-03 and RT-04 on the SH and GSH contents of ASMC.

Despite, different effects of the organotelluranes on GSH content of IRLM and ASMCs, loss of ΔΨ and Ca^2+^ homeostasis were observed for ASMCs (Figure 4). Figure 4 shows the effects of RT-03 on the cytosolic Ca^2+^ concentration (black line). The cytosolic Ca^2+^ concentration was labelled by Fura-2AM fluorescence that shifts the spectrum peak from 380 to 340 nm after Ca^2+^ binding. The 340/380 nm intensity ratio increased progressively to the maximum within 5 min in response to the increase of cytosolic Ca^2+^ concentration. In the following, cytosolic Ca^2+^ concentration decreased due to mitochondrial uptake and attained the minimal around 25 min after the organotellurane addition. In the same conditions, the mitochondrial transmembrane potential (ΔΨ) was measured for ASMC after the addition of the organotellurane. The loss of ΔΨ promoted by RT-03 was accompanied by decay of TMRM (gray line) fluorescence at 340 nm. The halftime (t_1_) of TMRM fluorescence decay was determined as 7.8 min by fitting the data using equation 2,

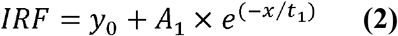

Where y_0_ is the offset, A_1_ the decay amplitude and t_1_, the halftime of the decay. The halftime of ΔΨ leakage was coincident with the maximum of the cytosolic Ca^2+^ overload and indicated by the vertical red dotted line in Figure 4. The cytosolic Ca^2+^ buffering promoted by the mitochondrial uptake contributes to the organelle permeabilization. The results obtained with RT-04 was very similar (not shown). The inset of Figure 4 shows snapshot of ASMC immediately and 13 min after the addition of RT-03 as indicated by the letters *a* and *b*. Loss of ΔΨ promoted by RT-03 and RT-04 was also observed using JC-1 as marker (not shown).

**Figure 4.**
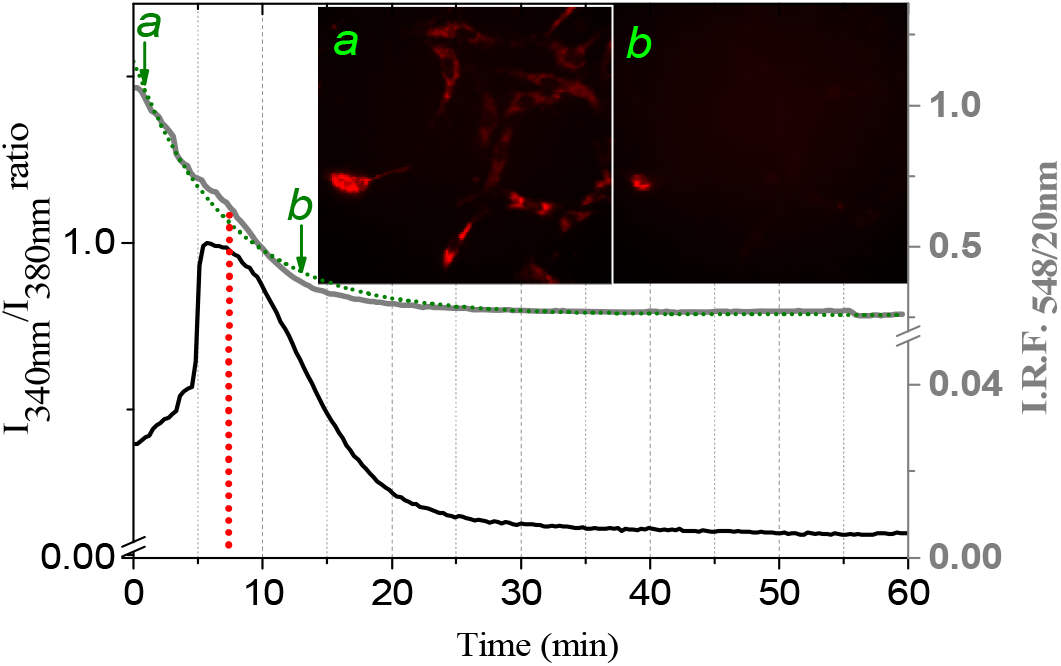
RT-03-promoted changes in Ca^2+^ homeostasis and mitochondrial permeabilization. ASM cells were grown on glass coverslips and incubated with TMRM and Fura-2AM in the presence of 50 μM of organotelluranes. The black line represents the 340/380 nm ratio of Fura-2AM fluorescence that increases proportionally to the cytosolic Ca^2+^ concentration. The gray line represents the ΔΨ dissipation curve that was fitted by equation 2 (green dotted line). The vertical red line marks the half-life of the ΔΨ decay that is coincident with the maximum of Ca^2+^ overload in cytosol. The inset shows the snapshot of ASMC labeled by TMRM at the times indicated by the green arrows.

Considering the effect of the organotelluranes on mitochondrial bioenergetic, it was investigated the capacity of RT-03 and RT-04 to promote the thiol depletion of isolated adenine nucleotide translocase was investigated (Figure 5). The organotelluranes decreased drastically the thiol content of isolated ANT that is consistent with mitochondria permeabilization promoted by these compounds.

**Figure 5.**
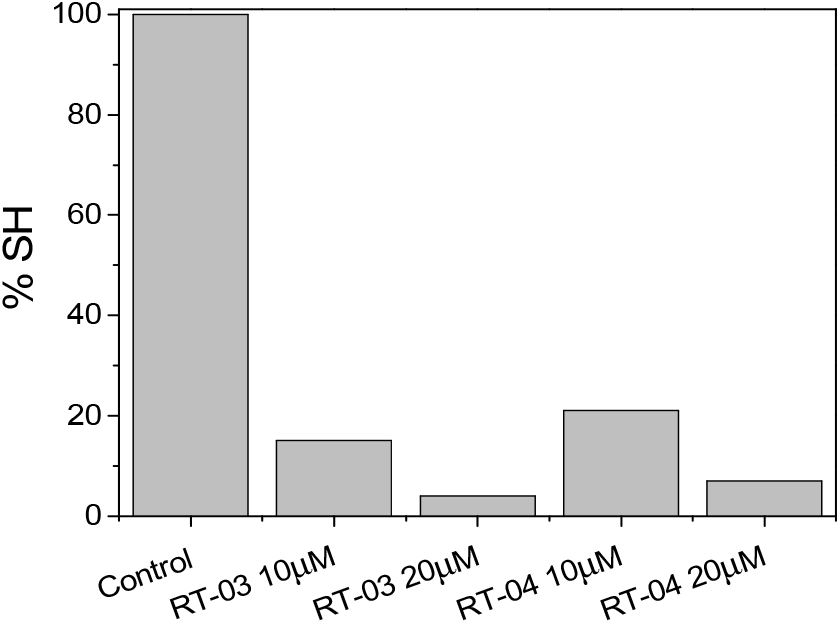
Effect of organotelluranes on the thiol content of ANT. ANT was isolated and incubated for 15 min in the presence of RT-03 e RT-04 and the thiol content determined colorimetrically at 412 nm after the reaction with DTNB.

The effects of organotelluranes (50 ◻M) on the ΔΨ, SH content and Ca^2+^ homeostasis of ASMC culminated with cell death that was evidenced by changes in the nuclear morphology (Figure 6).

**Figure 6.**
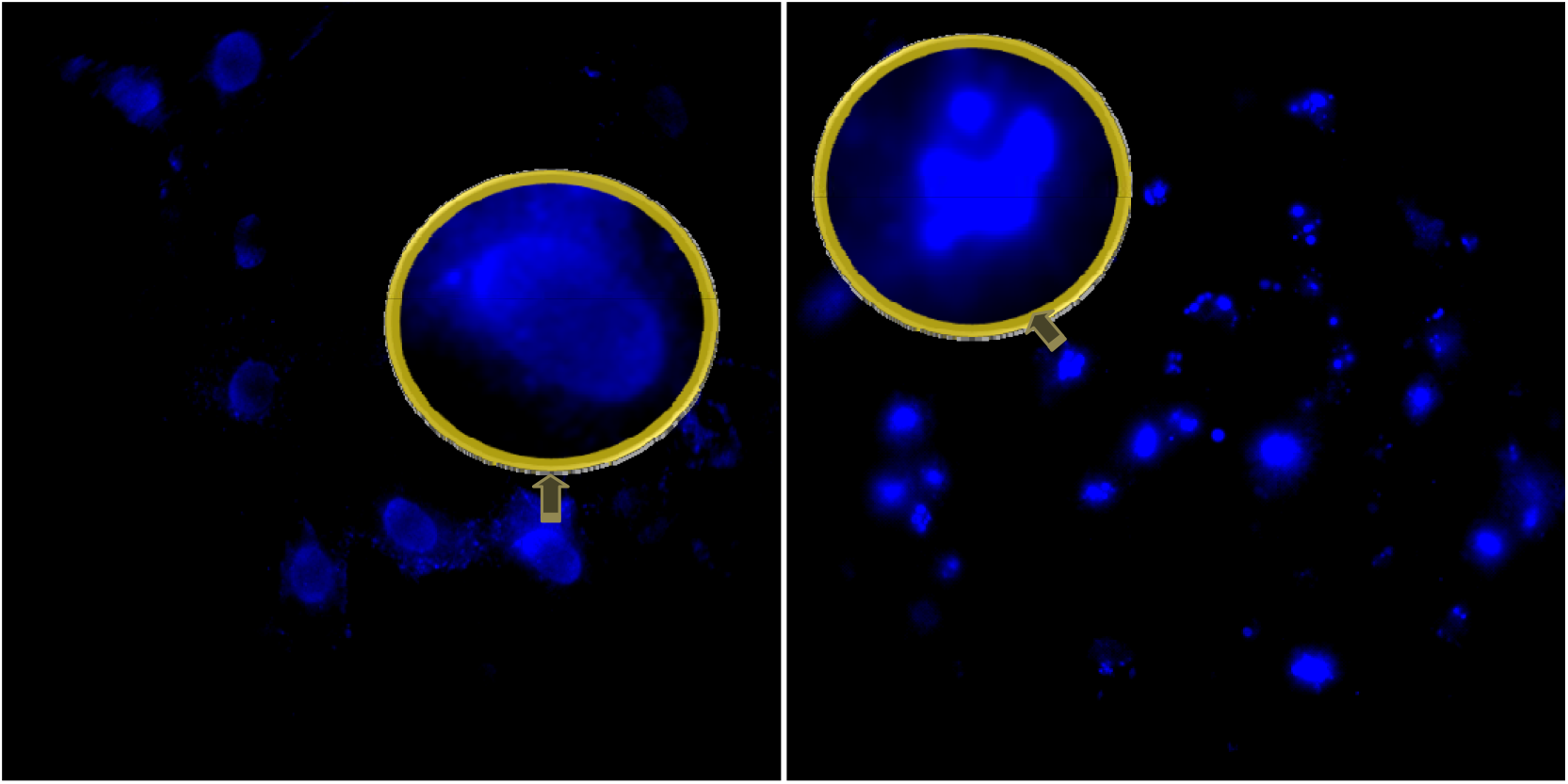
Effect of RT-03 on ASMC morphology. Snapshots show ASMCs nucleus after 15 h of incubation in the absence (left panel) and presence (right panel) of 50 μM of RT-03. The cells were incubated during 15 and the nucleus labeled with Hoeschst 33342.

Figure 6 shows the snapshots representative of cell morphology after 15 h of incubation with RT-03. After incubation, cells were treated with aqueous solution of Hoechst 33342 (1μg/ml), for 15 min. The images show the normal morphology of control cells (left panel) and the typical apoptotic nucleus with the characteristics of condensation and pycnoses. RT-03 promoted also loss of ASMC adherence and even the cells that remained stuck on the plaque presented pycnotic nucleus. In the following, the effect of RT-03 and RT-04 on the expression of proteins related to redox balance of cells was determined by real time PCR (Figure 7A-L).

**Figure 7.**
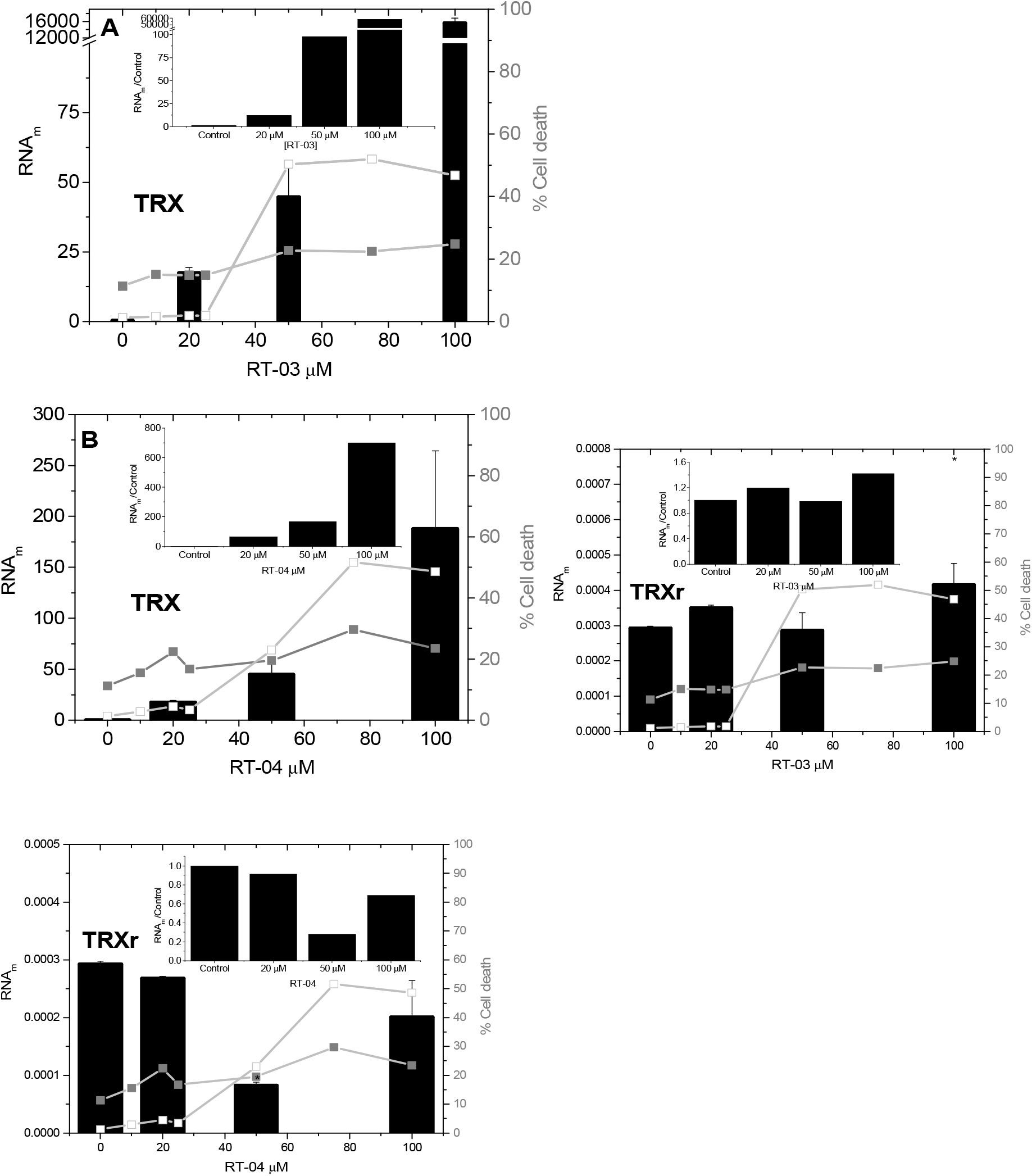
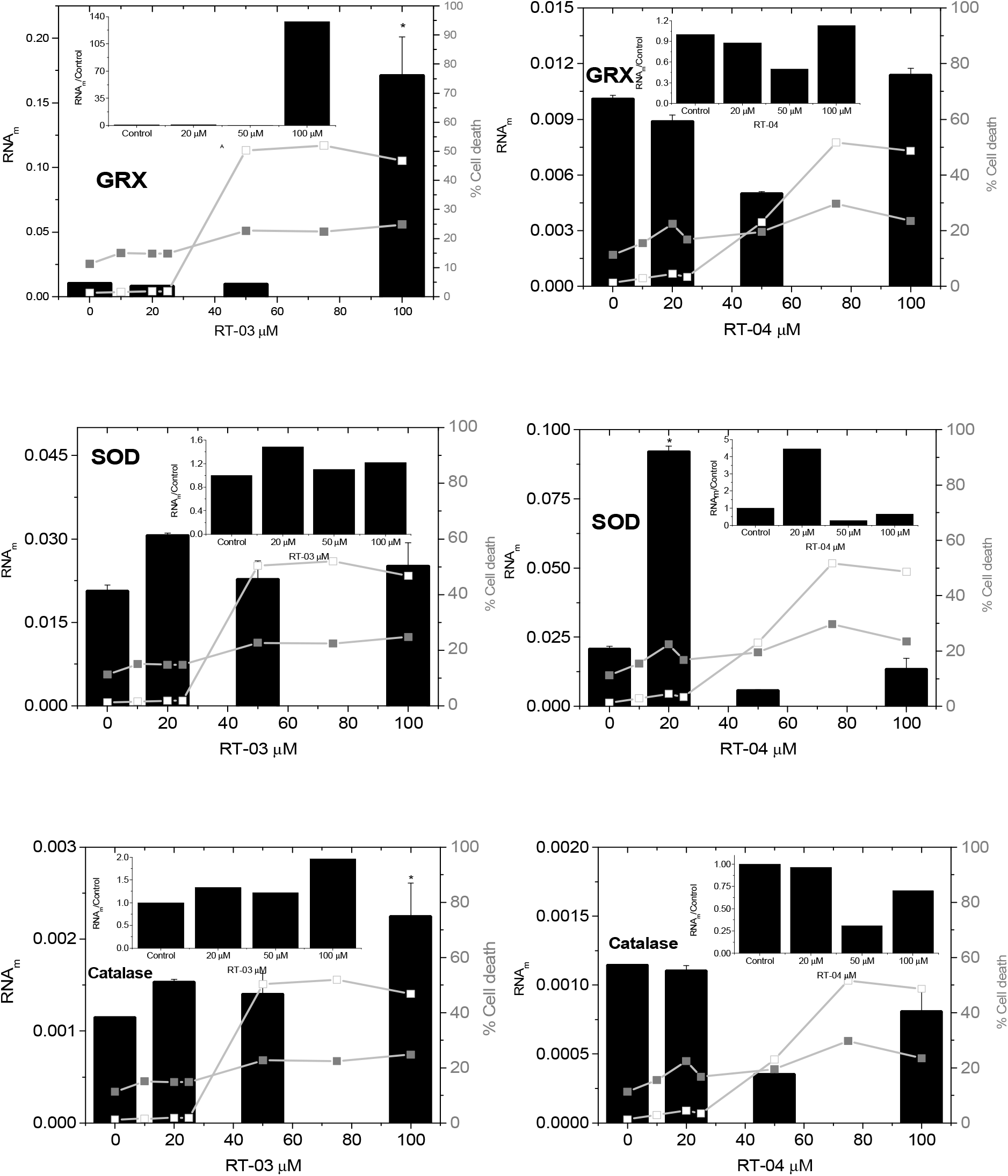
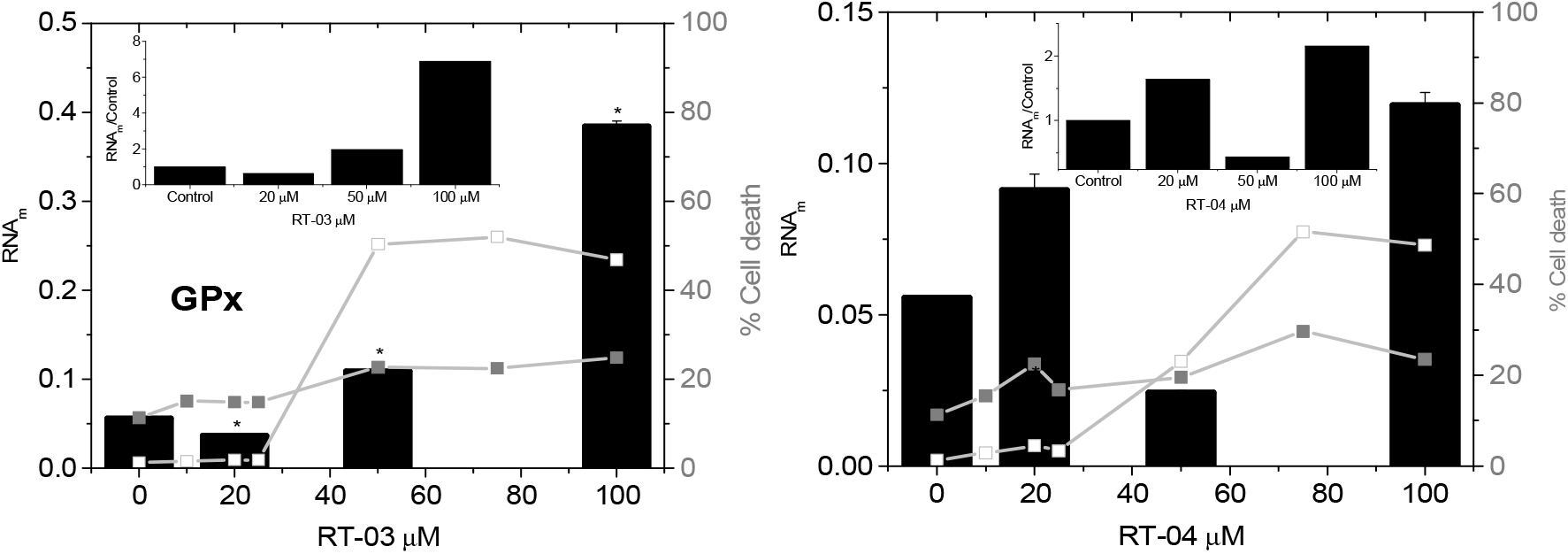
Effect of RT-03 and RT-04 concentrations on the cell death mechanisms and the expression of antioxidant enzymes.

In Figure 7A-L the data of RNAm production was shown overlapped with the percentage of apoptotic and necrotic ASMCs that were treated with the corresponding concentration of RT-03 and RT-04. Thioredoxin (TRX) had the expression extremely enhanced by the organotelluranes, particularly, RT-03. The results obtained with 100 μM of RT-03 suggests a total uncontrol of the TRX RNAm production. Contrarily, the organotellurane RT-03 did not affect significantly the expression of thioredoxin reductase (TRXr) while RT-04 promoted an inhibition its expression. In the case of GRX expression, only 100 μM of RT-03 promoted a significant increase of the RNAm production. The expression of superoxide dismutase (SOD) increased more significantly in ASMCs treated with 20 μM of organotelluranes and in this condition, the more severe effect was observed for RT-04. The concentration of 20 μM produced the greatest increase of apoptosis as the preponderant mechanism of cell death. In concentrations >20 μM, the expression of SOD was inhibited by both the organotelluranes, and more significantly by RT-04. The expression of catalase was also decreased significantly in ASMCs treated with 50 mM of RT-04 but not by RT-03 that duplicate the enzyme expression only at 100 μM. RT-03 also promoted increase of glutathione peroxidase (GPx) expression in a concentration-dependent manner. RT-04 also increased the expression of GPx, except at the concentration of 50 μM in which a significant decrease of the enzyme expression was observed.

## Discussion

Organotelluranes have a particular reactivity with the thiol groups herewith their biological activities are rationalized. The reaction of a thiol with organotelluranes is still a matter of debate since the rate of such processes difficult to access in experimental model systems; an early attempt was conducted by Albeck and coworkers who proposed two mechanistic pathways by which these reactions would occur, an associative and dissociative mechanisms (Scheme 1). (Cunha et al. 2009; Yokomizo et al. 2015)

**Scheme 1.**
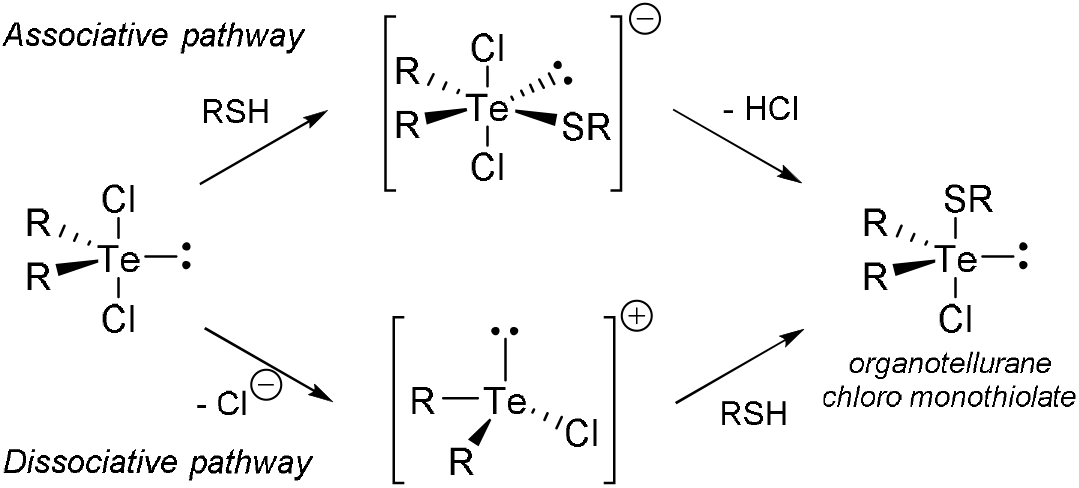
Proposed ligand exchange reactions of dichloro organotelluranes.

The difference of both mechanisms lies in the nature of the reaction intermediate as an anionic pentavalent intermediate is formed by addition of the nucleophile in the former mechanism whereas a cationic trivalent specie is formed in the later prior to the reaction with the thiol. Interestingly, orgatelluranes RT-04 and a dichloride derivative of RT-03 shows a remarkable stability in aqueous dimethylsulfoxide solutions, even in acidic media, showing that DMSO solutions of organotelluranes are very stable remaining in the form of dichlorides.(Princival et al. 2017) In general, the most commonly exploited organotelluranes in biological studies have a limited solubility in aqueous media thus theoretical studies have been carried out to investigate the fate of such species in water media since the hydrolysis can be anticipated. These studies indicate that after a series of ligand-exchange reactions, a thermodynamically stable telluroxide may be formed which is reactive towards thiols, nonetheless(Silva et al. unpublished data; Teixeira et al. 2018). The spontaneous formation of telluroxides from dihalo diorganotelluranes possessing an intramolecular coordination ether moiety further corroborated these theoretical studies.^92^ Telluroxides may likely react with thiols but through a nucleophilic addition mechanism forming an monothiolate that can react with another thiol leading to a dithiolate that can disproportionate forming a telluride and the disulfide (Scheme 2).

**Scheme 2.**
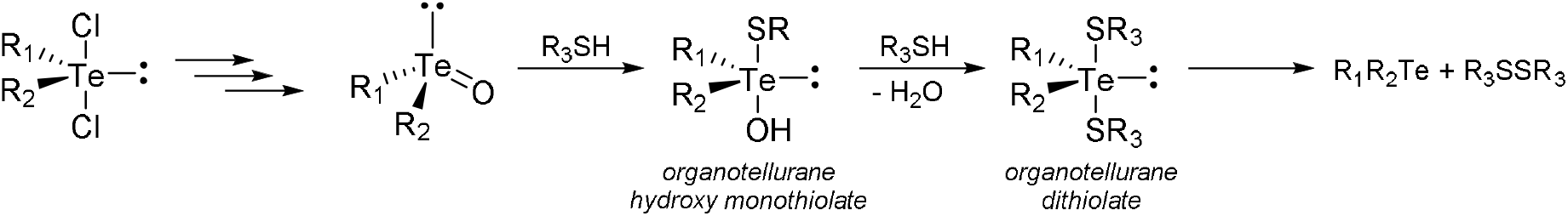
Reactions of thiols with organotelluroxides.

Organotellurane monothiolates originated in reactions with macromolecular thiols are likely to be kinetically more stable than those formed in reactions with low molecular weight thiols as cysteine, whose formation of the dithiol is more favorable due to lower steric hindrance. Whatever the mechanism, an intermediate protein-tellurane complex might form and become a target for the reaction with GSH and TRX. Also, the direct reaction of cysteine residues of TRX with organotelluranes cannot be discarded as it possesses more nucleophilic thiols than those of GSH and TRX. The protein-tellurane complexes can react with antioxidant proteins as TRX, GRX, and GSH by bis-thiolation leading to a redox pathway. In the redox pathway, glutathionylated protein or a mixed disulfide complex with TRX could be formed and subsequently recycled by TRXr and GRXr. However, inhibition of TRXr, previously described for some of organotelluranes, could prevent the recovery of protein thiol groups.(Albeck et al. 1998) In the case of TRX, cells responded to the impaired recycling of TRX with its overexpression. In this regard, RT-03 could more strongly inhibit TRXr and form more stable complexes with proteins leading to more drastic effects of TRX and GRX overexpression. The structure-dependent reactivity of RT-03-protein complexes via bis-thiolation (i.e., the formation of dithiolates) or redox mechanisms remains uncovered.

Before analyzing of the concentration-dependent effect of RT-03 and RT-04 on the expression of antioxidant enzymes, it is important to consider that for RT-03, but not for RT-04, the concentrations of 50 and 100 μM lead to a similar percentage of cell death. Also, RT-03 promoted overexpression of Grx, catalase, and GPx only at the concentration of 100 μM. Therefore, it is reasonable to consider that 100 μM of RT-03 promoted a severe disruption of cellular redox control and their effects are not comparative with the corresponding concentration of RT-04. Thioredoxin was the protein with the highest expression increase in response to treatment with RT-03 and RT-04. Overexpression of Trx was observed even below the EC_50_ of RT-03 and RT-04. Apart from the effect of 100 μM of RT-03, Grx was not overexpressed by ASMCs treated with 20 and 50 μM of RT-03 and had a decreased expression in the presence of 50 μM of RT-04. The expression of TrxR was not significantly affected by RT-03 and was inhibited by RT-04, more significantly above the EC_50_.

These results can be understood based on the following reasoning. The reaction mechanisms of telluranes with thiols have been extensively studied and can comprise two stages: the formation of the tellurium-sulfur bond forming a monothiolate and possible secondary reactions of the Te-protein complexes with nucleophiles. The formation of organotellurane adducts with a protein thiol group may occur through associative and dissociative mechanisms described previously. The protein complexed with the organotellurane may be the target of nucleophilic attack by glutathione, another protein, or another cysteine residue of its chain. Thus, the tellurane-protein adduct can be glutathionylated or form intra− or inter-molecular disulfides (RS-SR).

In any of these conditions, the protein may be recycled to its native form by the action of the Grx and Trx systems. Glutathionylated proteins can be reduced by the action o Grx and other disulfides by the Grx and Trx systems. Organotelluranes can inhibit TrxR impairing the recycling of oxidized Trx to the reduced form. The high consumption of reduced Trx without recycling by TrxR could lead cells to respond with the overexpression of the enzyme.

The absence of glutathione depletion suggests that the balance of oxidation and recycling of the peptide was not affected by RT-03 and RT-04 at the concentration range of 20-100 μM. Notwithstanding, GSH was not the preferential nucleophile that reacted with the Te-protein adducts. The reaction with GSH should be unfavorable because the pKa of its thiol group is 8.3, and most of the thiol is therefore protonated. Also, the organotelluranes are highly hydrophobic and should partition preferentially inside cell membranes. Taken together, these results suggest that membrane thiol groups are the principal nucleophiles reacting with RT-03 and RT-04 in ASMCs. Previously, it was demonstrated that RT-03 and RT-04 are also an efficient antioxidant for membrane lipids, as these organotelluranes did not promote organelle damage when used in low concentrations at nanomolar scale; however high concentrations of RT-03 and RT-04 promoted the opening of the MPTP associated with thiol depletion but without lipid oxidation in mitochondria.

Consistently, 20 and 50 μM of RT-03 did not significantly affect the expression of SOD and catalase. Above the EC_50_, RT-03 increased the expression of GPx. However, GPx-1 is a selenocysteine-containing enzyme, and its expression has unique forms of regulation that involves the trace mineral selenium. Therefore, it is not surprising that tellurium-containing compounds could affect the expression of GPx. Further, GPx controls the cellular levels of hydrogen peroxide and consequently the signal transduction involving redox-sensitive cysteines. Several proteins are redox regulated by TRX, such as NF-κB subunit p50, hypoxia-inducible factor 1-α, the tumor suppressor p53, c-Fos/c-Jun complexes, and others. Overexpression of thioredoxin in cancer cells is associated with apoptosis inhibition. In pancreatic cancer, the complexation of TRX with apoptosis signal-regulating kinase 1 (ASK1) inhibits apoptosis. TRX binds to the N-terminal noncatalytic region of ASK1 that encompasses the amino acid residues 1-655 leading to the enzyme inhibition. Consistently, necrosis rather than apoptosis contributed to cell death promoted by RT-03 and RT-04. Additionally, the cellular feedback described herein may play a role when other organotelluranes showed antitumor activity in a murine melanoma model using B16-F10 cells (Paschoalin et al., 2019).

## Conclusions

The organotelluranes RT-03 and RT-04 have peculiar effects on the redox balance by interacting specifically with thiol groups of proteins with concomitant protection of membrane lipids. The attack of protein thiol groups and particularly the capacity to inhibit TRXr affected mitochondrial bioenergetic and consistently lead to a significant increase in TRx, GRx, and glutathione expression levels peroxidase. Three levels of damage previously observed in RLM treated with the organotelluranes were reproduced in ASMCs. In RLM, RT-03 and RT-04 promoted mitochondrial transition pore opening at the concentration range of 5-10 mM, alterations in the bilayer fluidity, phosphorylation impairment, uncoupling, transmembrane potential disruption, and depletion of mitochondrial reduced thiol groups at the range of 15-30 mM promoted, and complete RLM respiratory inhibition at 100 mM. Consistently, ASMCs exhibited a maximum of a 2-fold increase in apoptosis rate after the treatment with 25 mM of the organotelluranes without the significant occurrence of necrosis that increased fifty times at the concentration of 75 mM of RT-03 and RT-04 and total uncontrol of the RNAm production at 100 mM. The damaging effects of RT-03 and RT-04, both in rat liver mitochondria and cells, are related to the reactivity with thiol groups of proteins. These findings are essential for comprehending how organotellurane affects cells, essential for further therapeutic applications of these compounds.

## Declarations

## Acknowledgments

Authors thank FAPESP, CNPq, CAPES, and the UFABC’s Strategic Research Units NBB and NuTS for the financial support and David da Mata Lopes for the technical assistance.

## Author Contributions

FSP and CHY performed the experiments, analyzed data, and designed the figures. RLORC synthetized and analyzed the studied organotelluranes. ILNC and RLORC designed the research project, and wrote the manuscript. All authors reviewed and approved the manuscript.

## Availability of data and material

As far as is reasonable, the authors will provide data as requested.

## Ethics approval

Not applicable.

## Consent to participate

Not applicable.

## Consent for publication

Not applicable.

## Code availability

Not applicable.

